# aradeepopsis: From images to phenotypic traits using deep transfer learning

**DOI:** 10.1101/2020.04.01.018192

**Authors:** Patrick Hüther, Niklas Schandry, Katharina Jandrasits, Ilja Bezrukov, Claude Becker

**Author notes:** These authors contributed equally (Patrick Hüther, Niklas Schandry).

## Abstract

Linking plant phenotype to genotype, i.e., identifying genetic determinants of phenotypic traits, is a common goal of both plant breeders and geneticists. While the ever-growing genomic resources and rapid decrease of sequencing costs have led to enormous amounts of genomic data, collecting phenotypic data for large numbers of plants remains a bottleneck. Many phenotyping strategies rely on imaging plants, which makes it necessary to extract phenotypic measurements from these images rapidly and robustly. Common image segmentation tools for plant phenotyping mostly rely on color information, which is error-prone when either background or plant color deviate from the underlying expectations. We have developed a versatile, fully open-source pipeline to extract phenotypic measurements from plant images in an unsupervised manner. ara**deep**opsis was built around the deep-learning model DeepLabV3+ that was re-trained for segmentation of *Arabidopsis thaliana* rosettes. It uses semantic segmentation to classify leaf tissue into up to three categories: healthy, anthocyanin-rich, and senescent. This makes ara**deep**opsis particularly powerful at quantitative phenotyping from early to late developmental stages, of mutants with aberrant leaf color and/or phenotype, and of plants growing in stressful conditions where leaf color may deviate from green. Using our tool on a panel of 210 natural Arabidopsis accessions, we were able to not only accurately segment images of phenotypically diverse genotypes but also to map known loci related to anthocyanin production and early necrosis using the ara**deep**opsis output in genome-wide association analyses. Our pipeline is able to handle images of diverse origins, image quality, and background composition, and could even accurately segment images of a distantly related Brassicaceae. Because it can be deployed on virtually any common operating system and is compatible with several high-performance computing environments, ara**deep**opsis can be used independently of bioinformatics expertise and computing resources. ara**deep**opsis is available at https://github.com/Gregor-Mendel-Institute/aradeepopsis.

## Introduction

### The phenotyping bottleneck in plant -omics studies

Over the last decades, molecular techniques have steadily increased in throughput while decreasing in cost. A prime example of this is nucleic acid sequencing, which has followed a trend analogous to Moore’s law in computer science. However, phenotyping methods, i.e., methods aimed at determining the physical shape of an organism and at measuring morphological parameters, have not kept up with this pace, which has caused a “phenotyping bottleneck” [1] in the design and execution of scientific studies. This phenotyping bottleneck constitutes a major challenge also in plant biology, due to two major issues. The first is data acquisition, which in most plant phenotyping scenarios is equivalent to acquiring standardized images of plants growing in a controlled environment. Plant phenotyping requires space and a dedicated infrastructure that can accommodate the developmental and growth transitions that occur over a plant’s life. Moreover, plant development needs to be phenotyped over relatively long periods of time. For example, in the case of the model plant *Arabidopsis thaliana*, a relatively small and fast-growing species, a phenotyping experiment typically runs for several weeks or months, depending on the phenotype of interest. In their vegetative phase, i.e., before they produce shoots, flowers, and seeds, Arabidopsis plants grow as relatively small, flat rosettes and can therefore be considered two-dimensional. Because the whole plant area is visible from the top during this phase, top-view phenotyping to determine size, growth rate, leaf development, etc. is straightforward. While this type of high-throughput image acquisition has become almost trivial, robustly and faithfully extracting meaningful data from these images has not.

Overcoming the data acquisition challenge, for example by means of a dedicated plant phenotyping facility, therefore does not mitigate the second cause of the phenotyping bottleneck, which is data analysis. In the context of image-based phenotyping data, this includes automated image processing for object detection, object segmentation, and extraction of quantitative phenotypic trait data.

### Challenges in automated image analysis

The first key step on the way from images to quantitative phenotype data is also the most difficult one: defining the area in the image that depicts the object of interest, in this case the plant. On a small set of images, this segmentation can be done manually by delineating the plant object using *ImageJ* [2,3] or similar software. On large image datasets with hundreds or thousands of individual images, which are typical for experiments from phenotyping platforms, this task must be automated, both to speed up the process and to neutralize user bias. Commonly used software for segmenting plant objects from digital images relies on color information. In simple terms, such color information is stored in tables, with information on each pixel stored in a corresponding cell. While grayscale images are stored in a single table, where each cell contains the grayscale value of the corresponding pixel, color image information is stored in several such tables. For images using the very common additive color model RGB, three separate tables store information on red (R), green (G), and blue (B) color channel intensities for each pixel. A very simple approach to differentiate plants from background is to assume that, since plants are green, one can assign all pixels that pass a certain threshold in the green channel to the object ‘plant’ and ignore all other pixels. This approach is called binary thresholding and works well if, and only if, the assumption is correct that all plants in the experiment are green and that the remaining area of the image is not. In reality, this assumption is often violated: plants can produce high quantities of anthocyanins under certain circumstances, which causes a red or purple hue; they can also develop chlorosis or become senescent, in which case they turn yellow or brown and, ultimately, white. Moreover, as they grow larger over the course of an experiment, the leaves of neighboring plants can protrude into the image area of the monitored plant, thus creating additional green areas that should not be taken into account. Even the background color can fluctuate, either because of constraints in the experimental setup, or because other organisms, such as algae, start growing on the soil surface. In summary, color-based segmentation is sensitive to image parameters that are difficult to control in a biological experiment. It therefore often needs to be verified by laborious and time-consuming visual inspection of the individual images to identify and correct erroneous segmentation and artifacts. We therefore sought to develop an alternative, more robust approach to image analysis.

### Machine learning methods in image analysis

Alternative approaches to object segmentation that solve the aforementioned problems of color-based segmentation are available. These are based on machine learning methods and include Gaussian mixture models (GMM) [4] or Naive Bayes classifiers [5]. While these approaches are highly flexible, their implementation on new datasets requires substantial programming knowledge.

More recently, successful alternative approaches to segment particular classes of objects from images have come from the deep learning field. Convolutional Neural Networks (CNNs) have proven invaluable for image classification and segmentation tasks [6–9]. They perform well on data that has a local structure, which is inherently found in images where values of neighboring pixels tend to be highly correlated and contain recurring structures such as corners and edges. These models are supervised, meaning that they are provided with a set of manually annotated images containing the “ground truth” for each pixel, from which the model will attempt to derive rules for the classification of pixels based on the training dataset. This training process is iterative and usually performed over many thousands of iterations, during which the model attempts to map input images to their corresponding ground truth annotations. In brief, during training, a loss function is calculated to estimate the error between input and output, and the model subsequently tries to minimize this error via back-propagation [10]. Weight matrices throughout the network layers are updated along a gradient, following the chain rule to calculate partial derivatives of the loss function with regard to the layer weights. Due to the non-convex nature of the loss function, error minimization typically reaches only local minima, requiring careful selection of model parameters and a large and diverse set of training data to avoid overfitting. The latter is usually scarce because ground truth data for supervised learning have to be generated by meticulous manual annotation. Depending on the nature of the desired feature to be extracted, such a task is typically labor-intensive and time-consuming due to the large number of data points required to train a well-generalizing deep learning model *de novo*.

However, in a process referred to as transfer learning, trained model architectures that do well at pixel classification can be re-trained on new data that contain new classes while retaining already trained weights in layers that extract low-level features such as shapes and edges. Consequently, to re-train a model on, for example, plant rosettes, the dataset on which the model was originally trained does not need to contain plant rosettes. Transfer learning is mainly about updating the last layer of the model, which is much faster and requires considerably less training data than designing and training a completely new model [11].

Here, we introduce ara**deep**opsis, a pipeline centered around the deep-learning model DeepLabV3+, re-trained for segmentation of Arabidopsis rosettes. Initially designed to simply identify image areas containing plants (rosettes), we further developed the tool to discriminate three different health states of leaves. We show how ara**deep**opsis can be applied to reliably segment Arabidopsis rosettes independent of their shape, age, and health state, which was not possible with color-based segmentation algorithms. By non-invasively extracting color index data from the segmented leaf area, our tool delivered highly resolved quantitative data that we successfully applied in genome-wide association (GWA) analysis of anthocyanin content. To the best of our knowledge, ara**deep**opsis is the first tool to return quantitative data on plant senescence, by which we were able to identify genetic variants that drive premature senescence. Because Arabidopsis is the most widely used model species in molecular plant biology and genetics, we centered our pipeline on this species and show that it can accurately segment other species, too.

ara**deep**opsis is a versatile and robust tool that can extract various biologically relevant plant phenotypes with unprecedented accuracy. Because the pipeline is written in *Nextflow*, researchers with little computational background can install and run it on their personal computer or in a high-performance computing environment.

## Results

### Color-guided segmentation can be misguided by image and object parameters

We wanted to explore the performance and accuracy of classical color-based segmentation and of a self-learning algorithm in segmenting Arabidopsis rosettes from a large, automated phenotyping experiment. We monitored growth of 210 Arabidopsis accessions from the 1001 genomes panel [12] in natural soil, under climate controlled conditions, with six replicates per genotype, from seed to flowering stage. Using an automated phenotyping system (https://www.viennabiocenter.org/facilities/plant-sciences/phenotron/), we recorded top-view images twice per day, cropped them to frames containing single pots, then subjected them to rosette area measurements. We first used *LemnaGrid* (Lemnatec), which relies on color channel information to segment plants in top-view images and is commonly used for Arabidopsis phenotyping [13]. This resulted in accurate segmentation of young and healthy plants composed mainly of green tissue (Fig 1). However, because the plants in our dataset were grown for long periods in natural soil, many accumulated high levels of anthocyanins in their leaves, either because of genetic determinants or as a general stress response to the natural soil and its abiotic or biotic composition. Others showed an onset of senescence at later stages of the experiment. On images of such plants, segmentation often failed completely or resulted in only partial or inaccurate segmentation of the plant (Fig 1), showing that color-based segmentation is sensitive to deviations from the expected color composition of the object to be detected.

**Fig 1.**
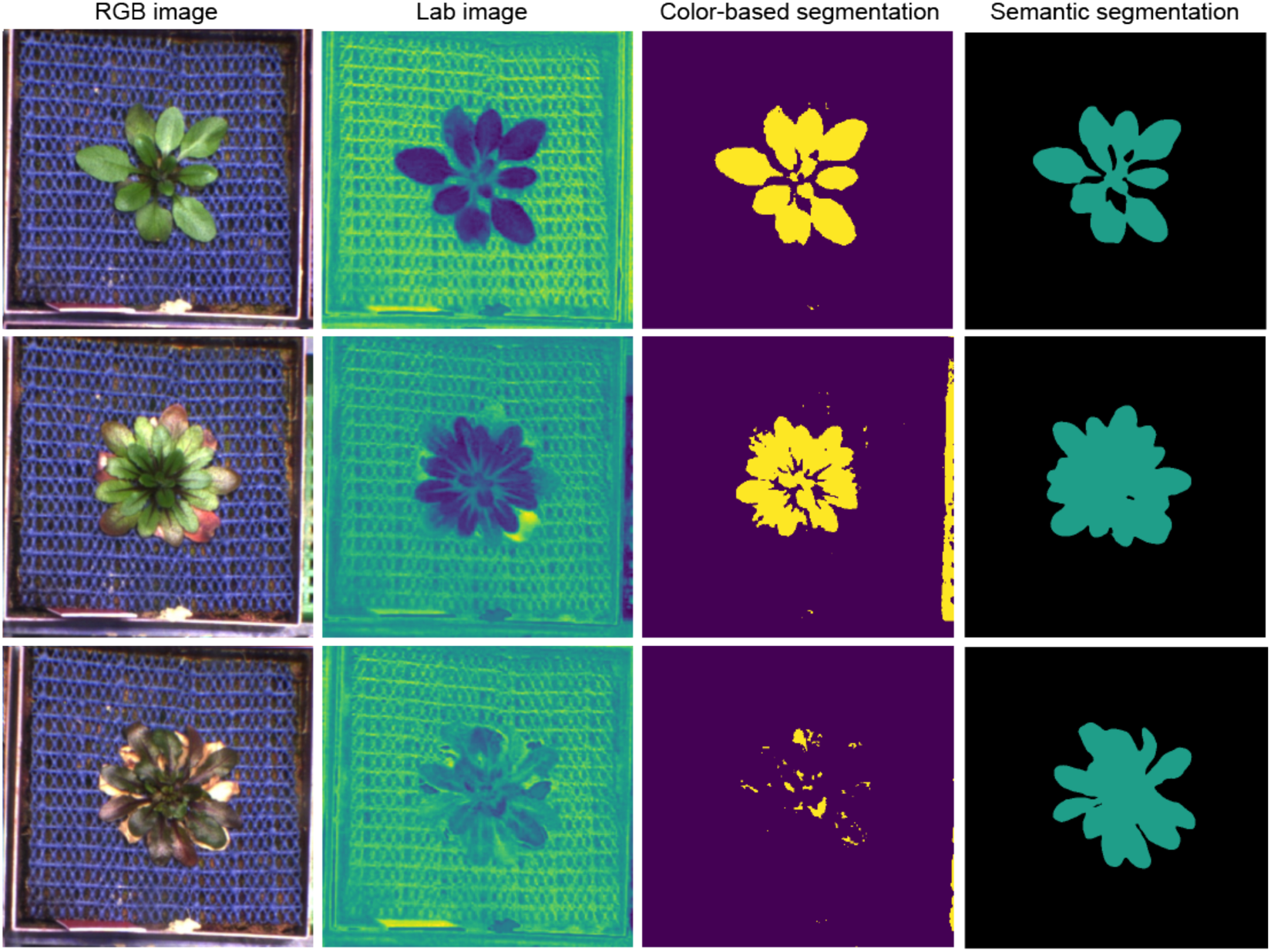
Performance of color-based vs. semantic segmentation. The leftmost column shows the original RGB images of a representative Arabidopsis plant from our phenotyping experiment at three different developmental stages. The second column shows the same images transformed into the Lab color space, which enhances contrast between green and other colors. Binary thresholding (third column), based on Lab input and used for color-based segmentation, results in difficulties to correctly segment older or anthocyanin-containing plants. Semantic segmentation by ara**deep**opsis is insensitive to color and contrast of the plant (rightmost column).

### Training of a proof-of-principle model

We hypothesized that self-learning algorithms would achieve higher accuracy while at the same time being more robust towards shifts in color patterns. We therefore developed ara**deep**opsis (*Ara*bidopsis *deep*-learning-based *op*timal *s*emantic *i*mage *s*egmentation), which is built around the deep-learning model DeepLabV3+. To test whether ara**deep**opsis could faithfully extract Arabidopsis rosettes from top-view images, independent of developmental stage or phenotype, we generated a ’ground truth’ training set: we initially generated a small dataset of 300 images, in which we manually annotated the rosette area, deliberately excluding senescent leaves. From these 300 annotations, we randomly selected 80% for model training and kept 20% separate for evaluating its accuracy.

After training, this model (referred to hereafter as model A) performed well on plants at various developmental stages and of different leaf coloring (Fig 1), which encouraged us to generate a much larger ground-truth dataset for more fine-grained models.

### Advanced models for differentiated plant area classification to classify individual leaves

From the example images shown in Fig 1, it is apparent that plant health changes not only in the context of the whole rosette but also between individual leaves or parts of leaves, with some accumulating high anthocyanin levels and others entering senescence. We therefore asked if we could train a model that would be able to semantically extract such features from leaves and assign them to different classes, depending on the phenotypic appearance. Not only would this allow finer resolution when assessing the overall health of the plant but would also enable segmenting differently colored areas.

Reasoning that it should be possible to segment senescent leaves separately, we generated a second, larger, and more finely annotated training dataset. This second dataset consisted of 1,375 manually annotated top-view images of Arabidopsis plants from the phenotyping experiment described above, from which we again kept 20% separate for validation. The images were selected semi-randomly such that they covered all various developmental stages and included healthy-looking as well as stressed plants that exhibited altered color profiles due to anthocyanin accumulation or senescence. Instead of manually annotating the whole rosette, as we had done for the initial training set, we annotated all leaves of each rosette individually and manually assigned each leaf to one of three different classes, depending on its appearance:

- green
- anthocyanin-rich
- senescent or dead

From these annotations, we generated image masks for two additional models complementing the initial one-class model A. For the two-class model B, we classified senescent (class_senesc) vs. non-senescent leaves (class_norm), whereas the three-class model C was trained to differentiate between senescent (class_senesc), anthocyanin-rich (class_antho), and green (class_norm) areas (Fig 2A). We then used these three annotated sets of masks for transfer learning of a publicly available *xception65* [7] based DeepLabV3+ checkpoint that had been pre-trained on the ImageNet dataset (see Methods) [14,15].

**Fig 2.**
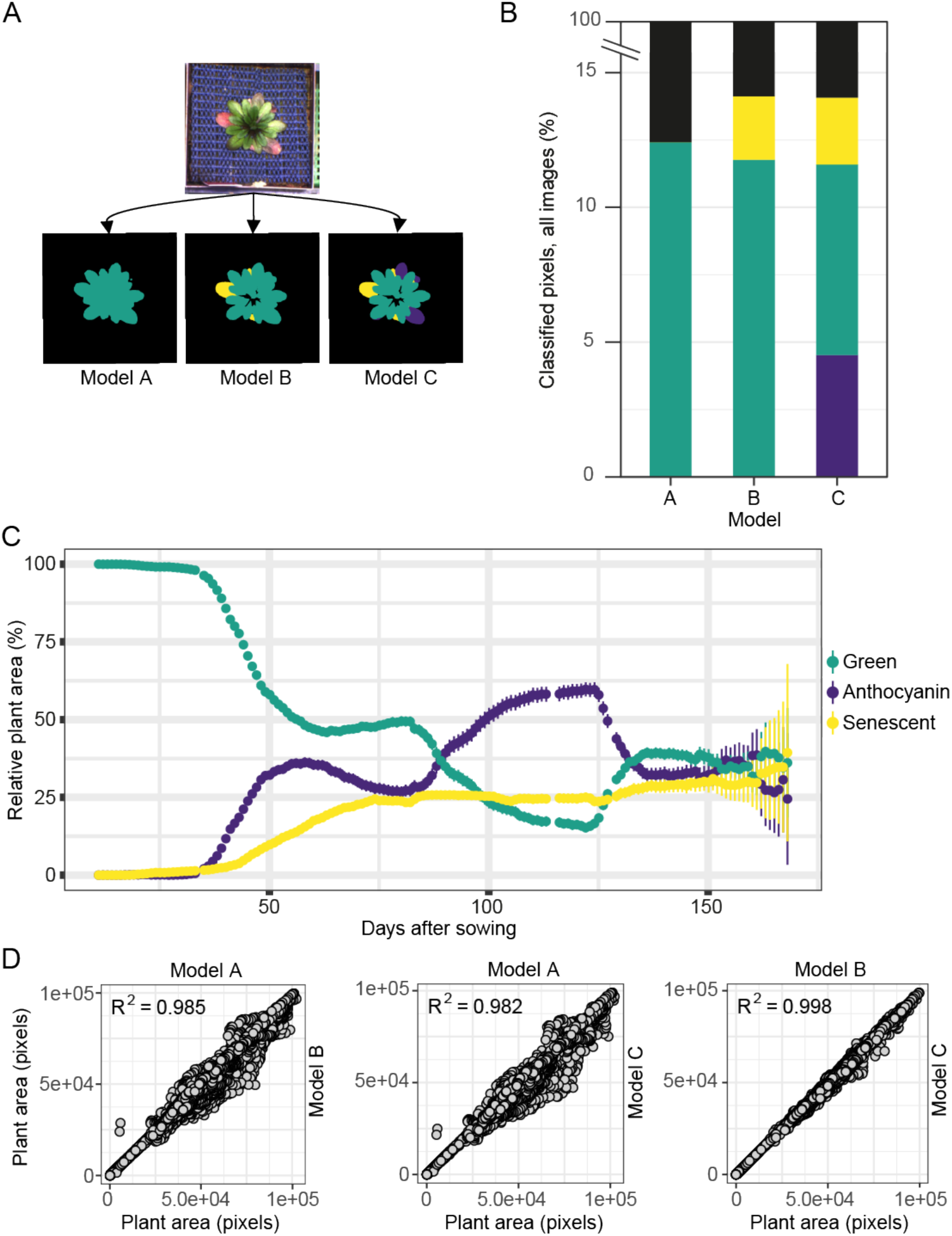
The three models available through ara**deep**opsis. (A) Example of segmentation results from the one-class model A, the two-class model B, and the three-class model C for a single Arabidopsis individual. (B) Overall comparison for percentage of all classified pixels across all 148,792 images between the three available models. (C) Relative number of pixels classified as green, anthocyanin-rich, or senescent by model C over time. The mean percentage of pixels assigned to each class is shown per day. Error bars indicate 95% confidence intervals. (D) Scatterplots showing pairwise correlations between measurements of plant area (includes all leaf states except senescence) between models A, B, and C.

After training of all three models had completed, we assessed their performance according to the mean intersection over union (mIoU) for all pixel-level classifications, which is defined as the mean fraction of true positives divided by the sum of true positives, false positives, and false negatives over all annotated classes. On the validation dataset, models A, B, and C reached an mIoU of 96.8%, 84.0%, and 83.1%, respectively.

Next, we compared how each model performed at segmenting all 148,792 rosette images of our dataset. First, we asked how much the different classes contributed to the area segmented as ’plant’. Averaged across all images, model A, which ignores senescent leaves, classified 12.5% of image area as plant tissue (Fig 2B). Models B and C both classified approximately 14% of image area as plant tissue (including senescent areas ignored by model A) (Fig 2B). The fraction of class_senesc segmentation was almost identical in both models. To further test whether our most complex model, model C, performed as expected, we analyzed the ratio of the three classes over the course of the experiment. As expected, we saw a sharp increase in anthocyanin-rich area beginning 30 days after sowing, followed by an increase in senescent segmentations 10 days later (Fig 2C). Relative classification of green tissue decreased accordingly. This showed that model C was able to capture an increase in senescence as plants grew older, accurately reflecting plant development.

Next, we compared whether the three models behaved differently in segmenting the combined rosette area. Using the total pixel area classified as leaves in each image, we found that models B (two-class) and C (three-class) correlated most strongly (R^2^ = 0.998; Fig 2D). Model A, our one-class model trained on annotations of whole rosettes and not of individual leaves, correlated less well with models B and C (R^2^ = 0.985 and 0.982, respectively) and had a tendency to segment larger areas than either of the other two (Fig 2D). Visual inspection of overlays of the segmentations produced by model A with the original image revealed that model A segmentations frequently included areas between leaves that were ignored by models B and C, explaining the larger rosette sizes measured by model A (see exemplary segmentations in Fig 2A). We believe that these differences are related to the different annotation strategies (whole rosettes in model A and more refined per-leaf annotations for the larger datasets in models B and C). When generating the ground truth for models B and C, we noticed that it is non-trivial to make consistent decisions as to when a leaf should be annotated as senescent or rich in anthocyanins, as the transitions are gradual, and classification can become subjective. This might also explain the decrease in mIoU scores that we observed with the two- and three-class models. We tried to mitigate user bias by having different individuals generate annotations, thus ensuring that the model would faithfully learn the features of interest.

### The ara**deep**opsis pipeline

The ara**deep**opsis pipeline is written in *Nextflow* [16]. The pipeline is fully open-source, licensed under GPLv3, and is presented in detail in the following sections. Briefly, the pipeline takes a folder of images as an input, splits the total image set into batches of equal size, and performs semantic segmentation on the images using a model of choice (see above “*Advanced models*…”). The semantic segmentation output is then used to extract morphometric and color index traits from each individual image (see below “*Extraction of morphometric and color index traits*”). Quality control and exploratory analyses can be carried out after the pipeline has finished by launching a bundled *Shiny* [17] application. In addition to offering some straightforward visualization of the results, *Shiny* also provides an interface to merge metadata and ara**deep**opsis output for downstream analysis (Fig 3).

**Fig 3.**
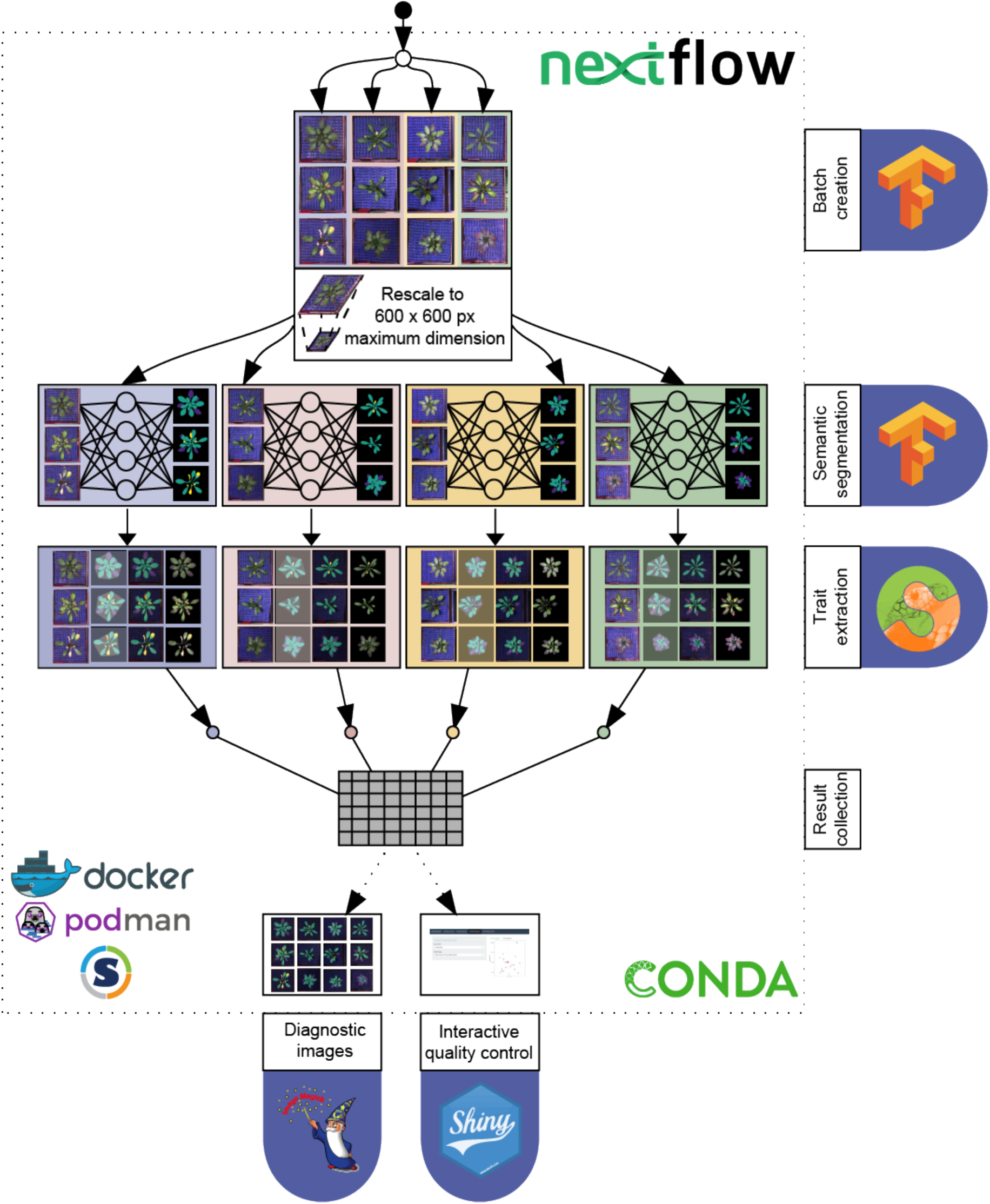
The ara**deep**opsis pipeline. A folder containing image files is passed to the pipeline. The total number of images is split into batches of a fixed size (indicated by different background colors). Batches are processed in parallel: first, the segmentation is performed on each batch, and then traits are extracted from the segmented images. The output from all batches is collected in one final table, the results table. In addition, the pipeline produces diagnostic images that show the segmentation results overlaid on the original image, color-coded segmentation masks, and background-subtracted plant rosettes. These diagnostic images can be explored in a *Shiny* [17] app, which is launched at the end of an ara**deep**opsis run, such that users can visually inspect the segmentations and verify their accuracy.

ara**deep**opsis is designed to segment plants from images that contain only a single plant, and we do not recommend usage on images containing multiple plants. In case the phenotyping system records whole or partial trays of plant pots, as is common for many automated phenotyping platforms, users will be required to pre-process these images and divide them into sub-images, each containing only a single individual plant.

### Extraction of morphometric and color index traits from segmented images

Next, we used the segmentation masks obtained from ara**deep**opsis to extract morphometric and color index traits of the plant (Fig 4), using the Python library *scikit-image* [18,19]). Besides the total number of pixels annotated as ’plant’, the pipeline returns the equivalent diameter, i.e., the diameter of a circle with the same number of pixels as the plant area. The major and minor axes of an object-surrounding ellipse are a measure of the aspect ratio of the plant and inform on whether the rosette is rather round or elongated along one axis. The area of a convex hull surrounding the object is a measure of plant size. The solidity of the plant object is calculated by dividing the area of the plant by the area of the convex hull. Finally, dividing the plant area by the area of its bounding box, i.e., a minimal box enclosing the object, is representative of the overall extent of the plant (Fig 4A). Depending on the type of downstream analyses, these indirect object parameters can be used as proxies for overall or specific growth.

**Fig 4:**
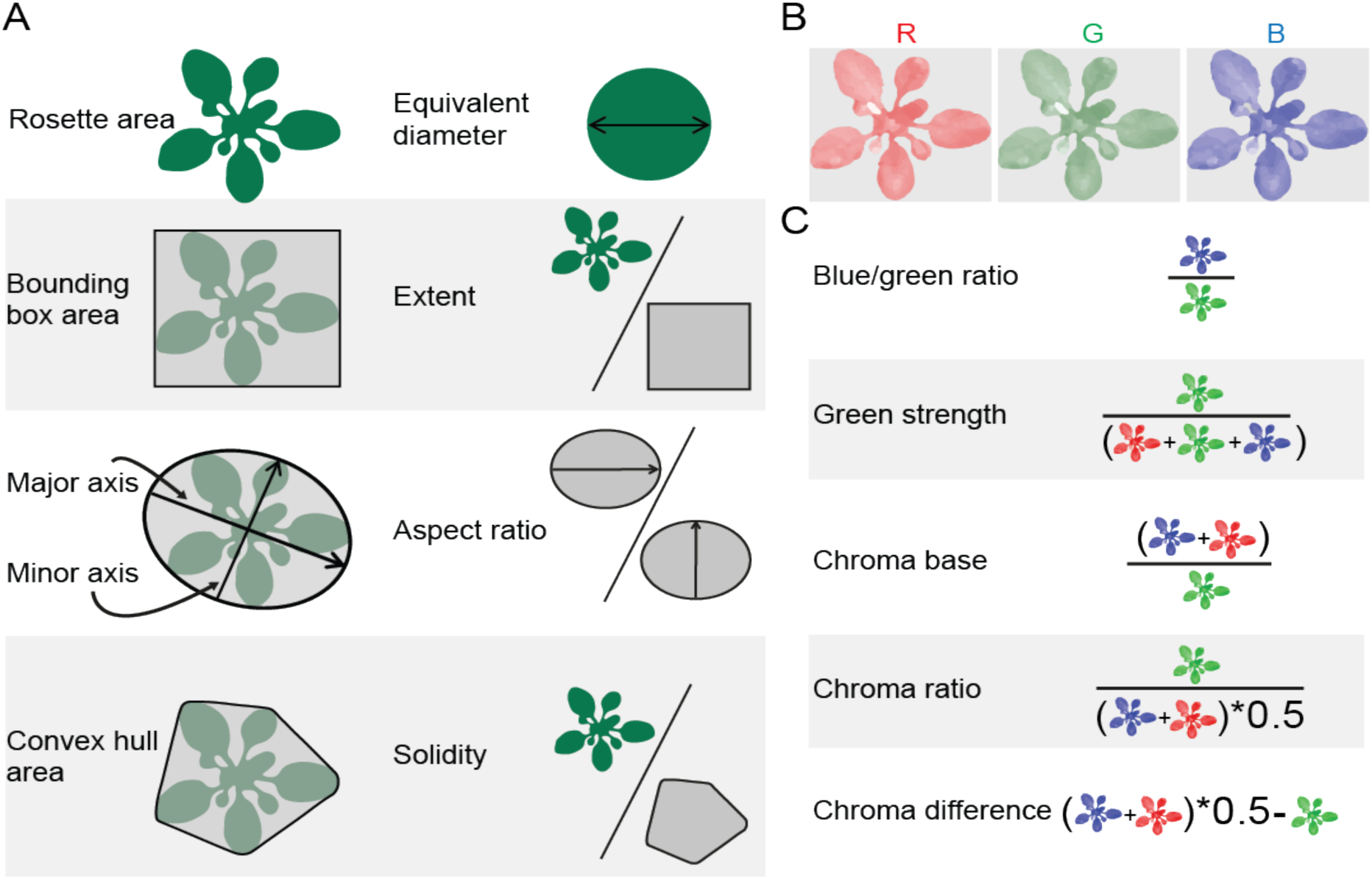
Morphometric and color index measurements extracted from segmentation masks. (A) Morphometric traits extracted using the Python library *scikit-image* [19]). (B) Separation of the segmented plant image into red (R), green (G), and blue (B) color channels. (C) Color channel indices calculated as described in [18]. The differently colored plants represent mean values of the different color channels; see (B).

The segmentation masks are also used to extract color index information from the plant area (Fig 4B). These traits are based on the intensity of each RGB color channel per pixel. Simple color index traits are, for example, the average intensity of the green channel over all pixels classified as ’plant’. We have implemented the color index traits described by [18] in our pipeline (Fig 4C). These traits are calculated for each class individually and for a compound class termed “plant_region”, which contains both class_norm and class_antho (only for model C). Details on the color index traits are shown in Fig 4C and provided in the Methods section.

### Validation against public datasets

To validate the accuracy of our pipeline, we reanalyzed publicly available images and compared the output of ara**deep**opsis to the published analyses. This served two purposes: first, we wanted to verify that the rosette features extracted by ara**deep**opsis segmentation were accurately reproducing published data. Second, we wanted to test whether our pipeline would remain robust when using data generated on different phenotyping platforms, with different image recording systems, pot sizes and shapes, and background composition. We used images from three published studies [20–22], generated on either the RAPA platform [23] at the Max Planck Institute for Developmental Biology in Tübingen, Germany [20,21], or the WIWAM platform (https://www.psb.ugent.be/phenotyping/pippa) at the VIB in Ghent, Belgium [22,24], and re-analyzed them using ara**deep**opsis.

Correlation analysis of the published rosette area derived from the RAPA system and our own measurements resulted in R^2^ values of 0.996 [21] and 0.979 [20], respectively (Fig 5A,B). Despite this overall very strong correlation, we noticed individual images in which our segmentation disagreed with the published one. When inspecting the respective images and segmentation masks from both analyses, we found that the respective rosette segmentations in the original analysis were inaccurate or incomplete, and that ara**deep**opsis segmentation was in strong agreement with the actual rosette in the image, even when plants had strong aberrant phenotypes or grew on low-contrast background (Fig 5A,B).

**Fig 5.**
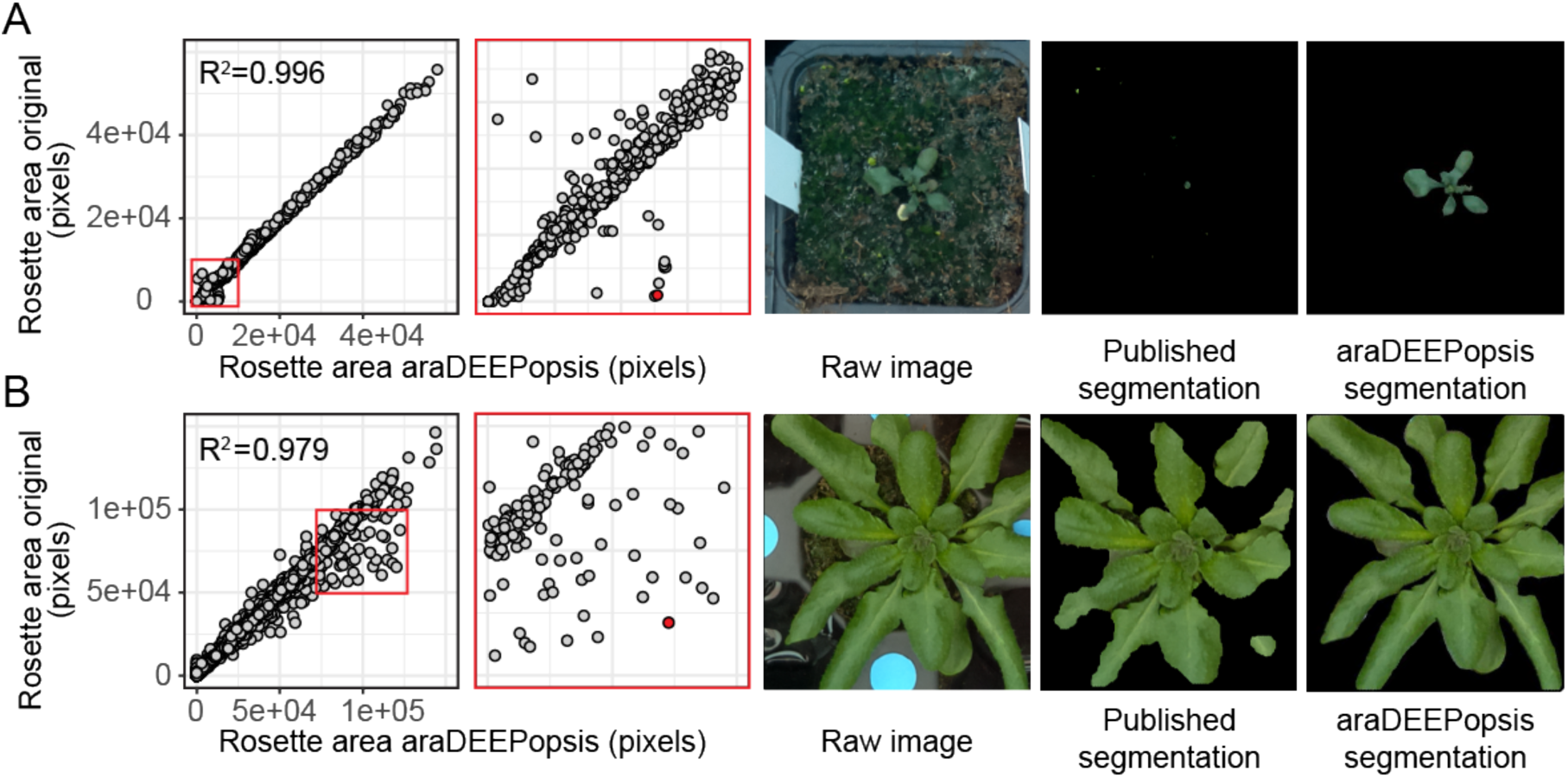
Validation of ara**deep**opsis output against published data. (A) Validation against data from [21]. The leftmost panel shows the correlation between values produced by ara**deep**opsis against published data; the boxed area is magnified in the second panel to highlight disagreements. The third panel shows the original image corresponding to the red data point in panel two. Panel four shows the original segmentation, panel five the segmentation by ara**deep**opsis. (B) Validation against [20] with the same order as in (A).

Re-analysis of the WIWAM dataset [22] also showed high correlation between the published measurements and our analysis (Fig S1). Unfortunately, original segmentations were not available for these images, but when inspecting some of the outliers, we again noticed highly accurate segmentation by ara**deep**opsis relative to the actual image (Fig S1).

### Validation by genome-wide association studies

#### Validation of color index traits

Often, the ultimate purpose of phenotype measurements is to relate them to genetic data in order to understand genetic determinants that control phenotypic traits. We therefore wanted to test whether ara**deep**opsis provided sufficiently high-quality data for Arabidopsis that could be used to search for genetic associations in genome-wide association (GWA) studies, relying on the 1001 genomes variant information [12]. Instead of morphometric traits such as size and growth rate, which are usually highly polygenic and therefore difficult to test, we wondered if we could make use of additional layers of image information provided by ara**deep**opsis as a proxy for physiological parameters. We hypothesized that RGB color information of the plant, i.e., the respective pixel intensities in the three color channels, would provide information on anthocyanin content and hence on the physiological state of the plant; it was previously shown that tissue color highly correlates with anthocyanin content [18]. Using the ara**deep**opsis segmentations, which accurately reflect the area covered by the rosette, we extracted color information by collecting data from the different RGB channels from the segmented object area, as explained above (Fig 4B,C). We then used *LIMIX* [25] to perform GWA analysis on the “chroma ratio” of the rosette area 37 days after sowing. The “chroma ratio” is calculated by dividing the mean intensities in the green channel by half of the sum of the mean intensities of blue and red channels (Fig 4C). It is therefore inversely correlated with the amount of anthocyanin accumulation and increases when anthocyanin content is low. We found strong associations with regions on chromosomes 1, 2, 4, and 5 (Fig 6A). When ranking by -log10(p), the second-highest ranking SNP is on chromosome 4 and the closest annotated gene from this SNP is *ANTHOCYANINLESS2* (*ANL2;* AT4G00730). *ANL2* encodes a homeobox transcription factor and has been implicated in controlling anthocyanin accumulation in sub-epidermal tissues [26]. Mutants in *ANL2* accumulate less anthocyanin in sub-epidermal leaf tissue. In our data, accessions carrying the alternative allele near *ANL2* displayed a higher chroma ratio, indicative of lower anthocyanin accumulation (Fig 6B). This shows that ara**deep**opsis is able to non-invasively extract quantitative color index data of the whole rosette that can be applied to genetic association analysis.

**Fig 6.**
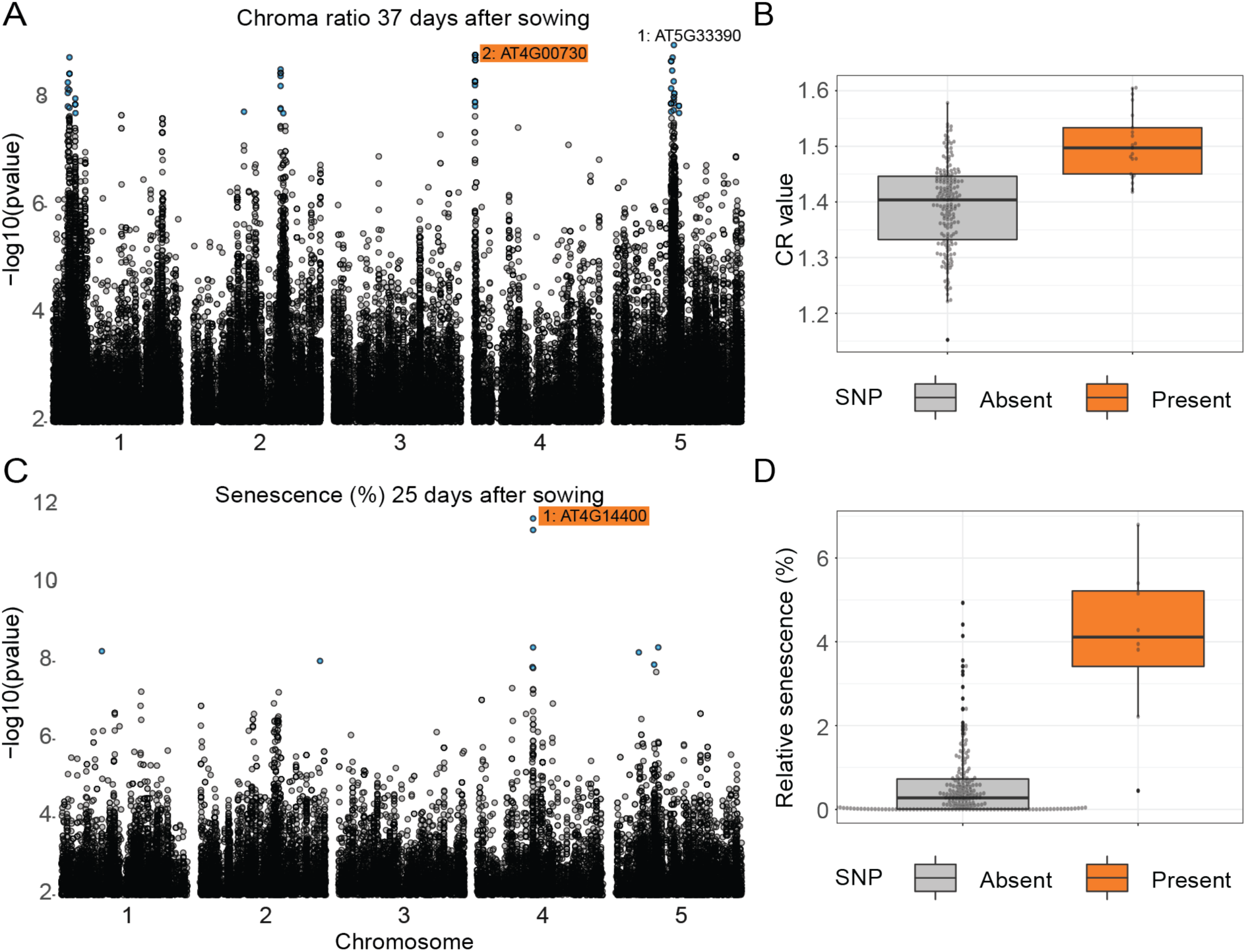
Genome-wide association (GWA) analyses based on ara**deep**opsis output. (A) Results of GWA on the trait “chroma ratio” 37 days after sowing. The log10-transformed *p*-value for each SNP is plotted (only SNPs with minor allele frequency >5% are shown). SNPs that are significant after Bonferroni correction are shown in blue. For the two highest-ranking SNPs, the closest gene is indicated. (B) Chroma ratio of plants 37 days after sowing, split by accessions carrying the reference and alternative alleles of the SNP close to *AT2G00730* (*ANL2*), highlighted in (A). (C) Same as (A) for the trait “relative senescence” 25 days after sowing. *AT4G14400* corresponds to *ACD6* (see text). (D) Relative senescence 25 days after sowing, split by accessions carrying the reference and alternative alleles of the SNP close to *ACD6*, highlighted in (C).

#### Validation of pixel-level classifications

Having validated our hypothesis that color profiles of the whole plant area can be used to extract informative traits, we asked if we would be able to identify genetic variants significantly associated with the relative amount of senescent or dead tissue during early development. We conducted a GWA analysis in which we used the relative amount of senescent tissue at 25 days after sowing as the phenotypic trait. This is early in the vegetative phase of plant development, and the relative senescent plant area is small compared to later stages (see Fig 2C). We found several genomic regions that displayed significant associations with this phenotype, the most striking of which was located on chromosome 4. The highest-ranking SNP in that locus was located within the coding region of ACCELERATED CELL DEATH 6 (*ACD6;* AT4G14400) (Fig 6C). The *ACD6* locus has been extensively studied in Arabidopsis and was identified as associated with vegetative growth, microbial resistance, and necrosis, in particular in F1 hybrids [27–29]. Our plants were grown on natural, non-sterilized soil from a field site near Zurich, Switzerland [30]; it is therefore fair to assume that the microbial load and potential biotic stress levels were higher than in standard potting soil. We observed an ~8-fold difference in median relative senescent tissue per plant (Fig 6D) from approximately 0.5% of total segmented pixels in accessions carrying the reference allele to around 4% in those carrying the alternative allele. This is in line with published studies, which showed that certain alleles of *ACD6* cause auto-immunity in Arabidopsis, which phenotypically manifests as premature senescence or necrosis [27].

### Analysis of other plant species

While our specific goal was to faithfully segment Arabidopsis rosettes, we wanted to see if our method would generalize to other plant species with similar overall morphology. We therefore tested ara**deep**opsis on top-view images of the Brassicaceae *Thlaspi arvense*, which has a similar leaf shape and phyllotaxis to Arabidopsis but does not form a flat rosette on the ground. Without training a model on images of *T. arvense*, i.e., using the model trained on Arabidopsis, segmentation masks matched the original plant highly accurately (Fig 7), showing that ara**deep**opsis is robust to variations in plant morphology, as long as these do not deviate from the general architecture that was used in the training set.

**Fig 7.**
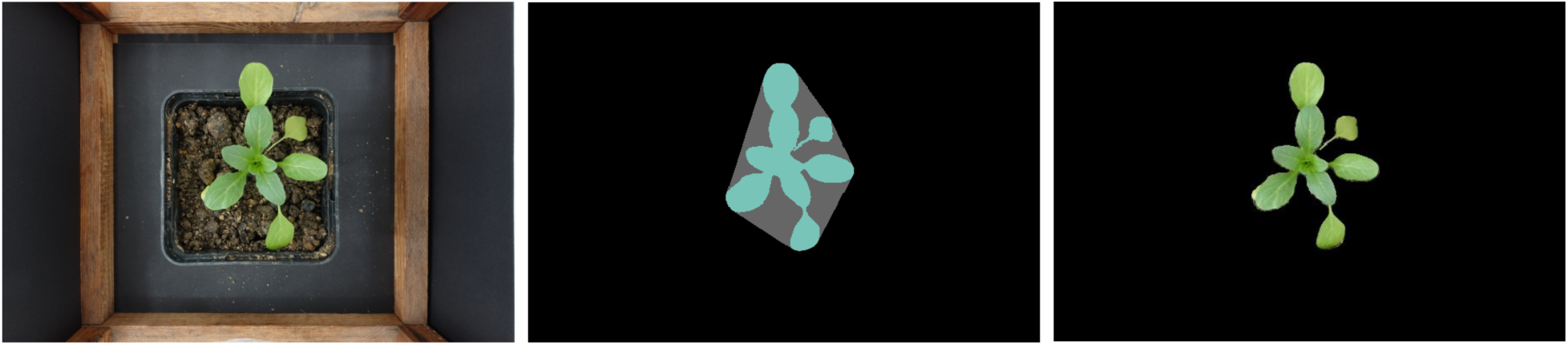
Segmentation of *T. arvense*. The leftmost panel shows the original image of a *T. arvense* individual. The middle panel shows the segmented mask and convex hull area. The rightmost panel shows the plant accurately cropped from the original image using the mask generated by ara**deep**opsis.

## Discussion

Here, we have presented ara**deep**opsis, a versatile tool to extract phenotypic measurements from small or large sets of plant images in an unsupervised manner. ara**deep**opsis is able to faithfully identify, segment, and measure plants from top-view images of rosettes, such as those of Arabidopsis, using deep learning methodology. Our tool is easy to use, runs on personal computers as well as high-performance computing environments, and is robust against variations in overall plant appearance, image quality, and background composition, making it superior to common color-segmentation based tools and accessible to a broad community of plant researchers.

### A fast and easy-to-use segmentation tool

Our models were trained on sets of 240 (model A) and >1000 (models B and C) images, respectively, with manually annotated ’ground truth’ segmentations, which required a non-negligible amount of manual labor. We believe that these investments were warranted, because ara**deep**opsis produced highly accurate segmentations and morphometric measurements that highly correlated with published data; occasional deviations from the originally reported data could in most cases be related to more accurate segmentation by ara**deep**opsis. Besides being accurate and requiring limited labor, the method is also fast: the fully automated analysis of 100 randomly selected images from our dataset took 12 minutes on a laptop computer (24 GB RAM, Intel i5-8250U 1.6 GHz). Depending on available resources such as memory and number of cores, the implementation in *Nextflow* enables straight-forward parallelization, allowing for a significant increase in speed when deployed in high-performance computing environments. At the same time, *Nextflow* is agnostic to the computational infrastructure being used, making it straightforward to deploy ara**deep**opsis on any type of computer. While training of the model greatly benefitted from the availability of GPUs, image predictions can also be carried out using CPUs in a time-efficient manner.

### aradeepopsis is robust to image characteristics and background composition

Many of the images in our training dataset had blue plastic meshes as background. This could raise the concern that the model might have learned to classify pixels belonging to the background rather than the plant of interest, i.e., to segment the plant as “non-background”, which would render the tool unreliable when using it on images with a different background composition. However, by testing ara**deep**opsis on images acquired in other phenotyping environments using different physical backgrounds, and on images of potted plants with substantial algal growth surrounding the Arabidopsis plant of interest, we observed that segmentation of leaves remained accurate and was agnostic to the background (see Fig 5A). Ideally, ara**deep**opsis should be used with images that are homogeneous within one set regarding light intensity, exposure time, etc. Segmentation still works for images with various light intensities (Fig S2), and plant size and geometry can still be analyzed, but quantification of color intensities based on the original image is no longer comparable in that case.

### Accurate determination of plant health state

ara**deep**opsis not only segments rosettes and extracts morphometric parameters, it also allows for the extraction of color channel information, which can be used to make assessments of the plant’s anthocyanin content and overall health status. We have shown that these results can be used, for example, for quantitative genetics approaches (Fig. 6).

Besides a simple one-class model (model A) that is able to segment non-senescent leaves, ara**deep**opsis includes two models that allow segmentation of anthocyanin-rich and/or senescent areas, depending on their color composition. As such, ara**deep**opsis is capable of reliably classifying senescent leaves and of distinguishing healthy, green leaves from stressed, anthocyanin-containing ones. Our models could also be extended to additional classes, depending on the specific needs of researchers and the phenotypes of interest.

## Outlook and Perspective

Our study shows that transfer learning is a valuable approach to overcome the phenotyping bottleneck. Our trained models have restrictions with regard to the angle at which the images are recorded and currently perform best for top-view images. The model was designed for the analysis of Arabidopsis rosettes but was also able to segment images of other species. We believe that the approach could be extended to a broader range of species and also to other types of plant images, e.g. side-view angles. These adjustments would require expert annotation of the corresponding images but can be performed by an experienced researcher and trained helpers in a reasonable amount of time. Alternatively, we would like to propose that pre-existing annotations from different research groups and various species be collected into a centralized repository that could serve as a powerful resource for phenomics in plant science and breeding.

ImageNet, the dataset that was used to pretrain the baseline model we built upon, contains images of dogs, airplanes, bicycles, etc., but no detailed annotations of plant species. It is therefore remarkable that ImageNet pretrained models enable such high accuracy when retraining the last layer with a relatively small set of 240 manually annotated images. Ultimately, this highlights the potential such publicly available databases and models hold for research and suggests that we should exploit such resources more frequently.

While our current implementation focusses on large-scale image data from HT phenotyping platforms installed in controlled greenhouse environments, we envision the approach to also be beneficial for field phenotyping purposes. For example, choosing a smaller network backbone such as *MobileNetV3* [31] could enable field researchers to measure plant morphometry on the go with their smartphone and could thus facilitate data collection and interpretation in the field.

## Methods

### Plant growth

Plants were grown in long-day conditions (16 h light (21°C), 8 h dark (16°C), 140 µE/m^2^s) with 60% relative humidity and watered twice per week. Seeds were sown on September 20th 2018, and plants were imaged two times per day. Plants were monitored daily for emerging inflorescences and flowering accessions were removed from the growth chamber. Plants that had not flowered by December 11th 2018 were subjected to vernalization: the chamber was cooled to 4°C (ramped). During watering, the temperature was raised to 5°C. This program ended on January 21st 2019.

### Training & Validation

The *Computer Vision Annotation Tool* [32] was used for manual image annotation; custom scripts were used to produce annotation masks from the XML output. The publicly available *DeepLabV3+* [14,15] code was modified to enable model training on our own annotated training sets. The code used for training as well as download links for our annotated training datasets is available here: https://github.com/phue/models/tree/aradeepopsis_manuscript/research/deeplab.

For model evaluation, we split the annotated sets 80:20: 80% of the images were used to train the model and 20% for its evaluation.

A transfer learning strategy was employed by using a model checkpoint based on the *xception65* architecture [7] that has been pretrained on the *ImageNet* dataset [33]. Starting from that, training was implemented in an asynchronous manner using between-graph replication on our in-house *Slurm* cluster, allowing the use of 16 Nvidia Tesla V100 GPUs across 4 compute nodes. The training was performed according to published protocols [14,15] with the following changes: to account for the number of GPUs, the training batch size was set to 128 and a base learning rate of 0.1 was applied and decayed according to a polynomial function after a burn-in period of 2,000 training steps. Images were randomly cropped to 321×321 pixels, and training was stopped after 75,000 iterations.

### Implementation

Based on the trained models, an image analysis pipeline was implemented in *Nextflow* [16] and is available at https://github.com/Gregor-Mendel-Institute/aradeepopsis. The workflow is outlined in Fig 3. *Nextflow* allows external pipeline dependencies to be packaged in a *Docker* container, which we provide at https://hub.docker.com/r/beckerlab/aradeepopsis/. The container can be run using *Docker* [34], *podman* [35], or *Singularity* [36]. Alternatively, dependencies can be automatically installed into a *Conda* [37] environment.

The pipeline follows a scatter-gather strategy to scatter the input data into equally sized batches of arbitrary size, allowing for parallel processing, after which the results are again collected into a single output table. After the initial splitting, all images in one batch are first converted to TFrecord files using *TensorFlow* [38]. The records are then served to the trained *TensorFlow* model of choice in order to generate segmentation masks, containing pixelwise classification.

In the next step, morphometric traits are extracted from the images using the *regionprops* function of the Python library *scikit-image* [19] on a per-class basis. Additionally, color channel information is extracted in the form of mean pixel intensities for each of the channels in the original RGB image. This happens within the region determined by the segmentation mask and is also done for each class. Based on the color channel means, chroma indices are calculated as follows [18]

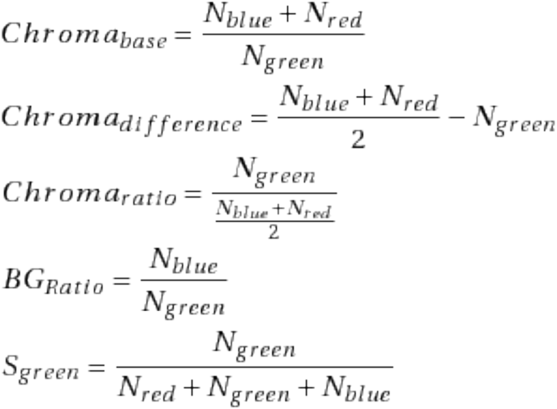

The results from these measurements are then collected from all processed batches, ultimately resulting in a single table containing a total of 78 traits. Based on the segmentation masks in combination with their corresponding input images, the pipeline also produces diagnostic images showing a color-coded mask, the cropped plant region, the convex hull, and an overlay image between the original and the mask.

These single-plant diagnostics are then, in an optional step, merged to produce summary diagnostics using *ImageMagick* [39], or can be viewed in an interactive *Shiny* application [17], allowing for fine-grained inspection of segmentation quality, pixel classification, correlations between traits, as well as time-resolved data visualization if appropriate metadata is provided.

### Genome-wide association studies

We performed GWA analysis on the traits produced by ara**deep**opsis using *LIMIX* [25]. We used the average of each trait per accession per day, and performed GWA analysis by fitting a linear mixed model using *LIMIX* [25]. To associate phenotype and single nucleotide polymorphisms, we used the 1,135 genotype SNP matrix and the corresponding kinship matrix, subset to those accessions where we had trait information. We screened the results for interesting trait-date combinations and followed these up using Arabidopsis-specific tools developed in-house (https://github.com/Gregor-Mendel-Institute/gwaR). The analysis is detailed in Supplemental File 1.

## Acknowledgements

We thank James M. Watson for critical reading of the manuscript and valuable comments. Karina Weiser Lobão helped in the manual image annotation. We thank Dario Galanti and Oliver Bossdorf for providing images of *T. arvense*. Klaus Schlaeppi and Selma Cadot provided the field soil. We thank Núria Serra Serra and Jorge Isaac Rodriguez for testing the analysis pipeline. The computational results presented were obtained using the CLIP cluster at VBC. Erich Birngruber gave advice on distributed model training on a Slurm cluster. We thank Stijn Dhondt for providing access to the PIPPA images via the BrAPI endpoint at VIB Ghent. Arabidopsis phenotyping was performed with support of the Plant Sciences Facility at Vienna BioCenter Core Facilities GmbH (VBCF), member of the Vienna BioCenter (VBC), Austria.

This work was supported by a grant of the European Research Council (ERC) to C.B. (grant ID 716823, “FEAR-SAP”), by the Austrian Academy of Sciences (ÖAW), and the Max Planck Society.

