## Supplemental Material for "aradeepopsis: From images to phenotypic traits using deep transfer learning"

SHORT TITLE:
Transfer learning in plant phenotyping

AUTHORS:
Patrick Hüther*, Niklas Schandry*, Katharina Jandrasits, Ilja Bezrukov, Claude Becker

#### Supplemental figures

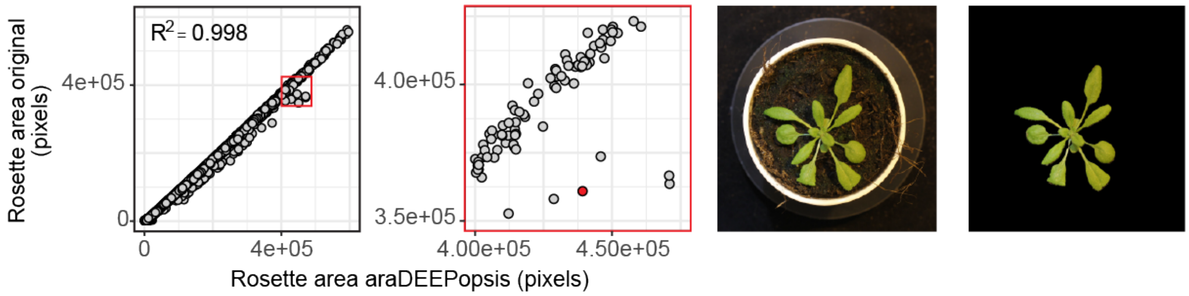


**Fig S1. Validation of morphometric measurements against published data from [22].** The leftmost panel shows the correlation between values produced by ARADEEPOPSIS against published data. The second panel is a magnification of the boxed area in the first panel, highlighting disagreeing measurements. The third panel shows the original image of the red data point in panel two. Panel four shows the segmentation by ARADEEPOPSIS which is in strong agreement with the original image, suggesting inferior segmentation in the published analysis.

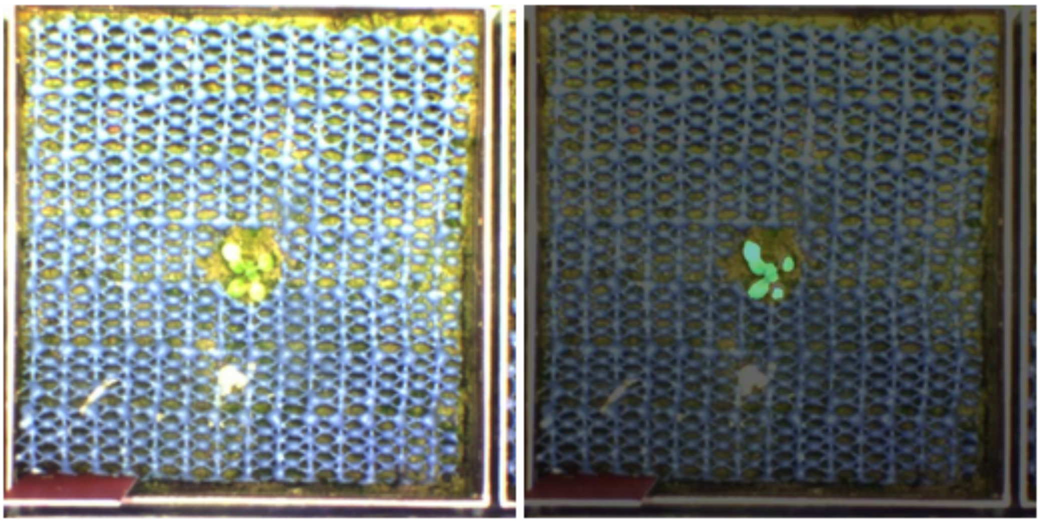

**Fig S2. ARADEEPOPSIS is tolerant to changes in image parameters.** Left: Original image, overexposed due to a technical problem with the light panel. Right: Segmented mask (green) overlayed with the original image.

### Supplemental files

**Supplemental File 1:** R-scripts used for the analysis of GWAS results.
https://syncandshare.lrz.de/getlink/fiLvKLBUsPKMrj4dLiGckiMa/araDeepopsis_supplement.html
